# Speed-dependent optimization of gravity effects for motor control

**DOI:** 10.1101/2023.03.14.532654

**Authors:** Gabriel Poirier, France Mourey, Cyril Sirandre, Charalambos Papaxanthis, Jeremie Gaveau

**Affiliations:** INSERM U1093-CAPS, Université Bourgogne Franche-Comté, UFR des Sciences du Sport, F-21000 Dijon, France

**Keywords:** Gravity, motor planning, effort minimization, optimal control

## Abstract

Several sensorimotor control studies have provided evidence supporting that the central nervous system optimizes gravity’s effects to minimize muscle effort. Recently, this hypothesis has been supported by the consistent observation of direction-specific negative epochs in the phasic electromyographic signal of antigravity muscles during vertical arm movements. This suggests that gravity torque is harvested to produce some of the arm’s motion. However, further investigation is needed to more finely understand how the CNS integrates gravity effects into muscle commands. Here, we aimed to analyze the phasic muscular activity across varying movement speeds during horizontal and vertical arm movements. We quantified the amount of negativity during acceleration and deceleration phases for all movement directions during fast, natural, and slow movements. We found that the negativity was more important during the acceleration phase of downward movements and during the deceleration phase of upward movements, resulting in diminished phasic activity compared to horizontal movements. Concomitantly, we found direction-specific effects of movement speed on phasic EMG activity of gravity muscles. This resulted in altered EMG to kinematics relationships in vertical movements compared to horizontal ones. These results support the *Effort-minimization hypothesis* and confirm that the negativity of phasic EMG is an important aspect of the motor command. Furthermore, the present results reveal that the CNS finely tunes this feature across a range of movement speeds and directions.

## Introduction

Due to its ubiquitous nature, the earth’s gravity acceleration plays an important role in sensorimotor control. Numerous researches have provided evidence that the central nervous system (CNS) has evolved an internal representation of gravity (Papaxanthis et al., 1998; Merfeld et al., 1999; Mcintyre et al., 2001; Indovina et al., 2005; Crevecoeur et al., 2009; Gaveau and Papaxanthis, 2011; Laurens et al., 2013b, 2013a; Gaveau et al., 2016). Although it was first hypothesized that the CNS uses this internal model to predict and compensate gravity’s mechanical effects on the body (the *Compensation Hypothesis;* Hollerbach and Flash, 1982; Atkeson and Hollerbach, 1985; Flanders and Herrmann, 1992), accumulative evidence support an alternative hypothesis according to which the CNS instead optimizes gravity’s effects to minimize muscle effort (the *Effort-Optimisation Hypothesis;* Papaxanthis et al., 2005; Lyons et al., 2006; Berret et al., 2008; Crevecoeur et al., 2009; Gaveau et al., 2011, 2014, 2016, 2021).

Consistent reports of direction-dependent arm kinematics indeed falsifies the *Compensation Hypothesis* whose predictions are independent of movements direction (Gentili et al., 2007; Le Seac’h and McIntyre, 2007; Gaveau et al., 2011; Bringoux et al., 2012; Gaveau et al., 2014; Yamamoto and Kushiro, 2014; Gaveau et al., 2016; Hondzinski et al., 2016; Rousseau et al., 2016; Yamamoto et al., 2016, 2019; Poirier et al., 2020; Gaveau et al., 2021). Numerical simulations have explained direction-dependent arm kinematics - and their progressive disappearance during microgravity exposure (Papaxanthis et al., 2005; Gaveau et al., 2016) - as the signature of a gravity-related optimization process that minimizes muscle effort (*Effort-Optimisation Hypothesis*; Berret et al., 2008; Crevecoeur et al., 2009; Gaveau et al., 2014; 2016; 2021). According to the *Effort-Optimisation Hypothesis*, the internal model of gravity would allow predicting and taking advantage of its effects rather than compensating for them.

According to the *Compensation Hypothesis*, the muscular command would be the addition of two components: the tonic component – that compensates for gravity – and the phasic component – that produces the acceleration (Flanders and Herrmann, 1992; Buneo et al., 1994; Flanders et al., 1994; D’Avella et al., 2008; Olesh et al., 2017). Recently, Gaveau et al. (2021) analyzed the phasic component of muscular activities during single degree of freedom vertical and horizontal movements. The authors observed consistent and direction-specific negative periods in the phasic activity of antigravity muscles during vertical movements. As predicted by the *Smooth-Effort Model* (Berret et al., 2008; Gaveau et al., 2014, 2016), those negative epochs appeared during the acceleration phase of downward movements and during the deceleration phase of upward movements. These results clearly support the *Effort-Optimization Hypothesis*, as negative epochs on phasic muscle commands mean that the hypothesized compensatory activity was lacking, thus that gravity torque produced some of the arm’s motion to accelerate and decelerate downward and upward movements respectively.

The analysis of muscle activation patterns provides a novel and strong support for the *Effort-Optimization Hypothesis* (Gaveau et al., 2021). Nonetheless, further quantifications of muscular commands are needed to more stringently test this hypothesis and to more finely understand how the CNS integrates gravity effects into muscle commands. One of the most basic and widely used descriptor of muscle commands is the tri-phasic burst pattern. Decades of research have consistently shown that, during point-to-point movements, muscle activations follow a tri-phasic temporal organization (Corcos et al., 1989; Cooke and Brown, 1994; Berardelli et al., 1996; Chiovetto et al., 2010; Forgaard et al., 2013; Huang et al., 2015; David et al., 2016). A first agonist burst produces the body-limb acceleration. Then, an antagonist burst produces its deceleration. Last, a second agonist burst stabilizes the final body-limb position (for a review, see Berardelli et al. 1996). This organization is so basic and stereotypical that its modification is used to underlie the pathophysiology of several neurological conditions such as Parkinson’s disease (Berardelli et al., 2001; David et al., 2016), Huntington’s disease (Thompson et al., 1988; Berardelli et al., 1999), dystonia (Van der Kamp et al., 1989; Münchau et al., 2001) or cerebellar ataxia (Hallett et al., 1991; Köster et al., 2002). Hitherto, most studies have investigated tri-phasic muscle patterns during movements performed in a transverse plane; i.e. during horizontal movements. Quantitative analyses have demonstrated that the first agonist burst is proportional to the movement acceleration peak, while the antagonist burst is proportional to the movement deceleration peak (positives correlations between EMG measures and kinematics; Corcos et al., 1989; Gottlieb et al., 1989; Cooke and Brown, 1994). According to the *Effort-Optimization Hypothesis*, gravity assists muscle force for accelerating (during downward movements) and decelerating (during upward movements) body-limbs. Thus, a straightforward prediction is that the well-known proportional relationship between muscle activity and arm kinematics will be altered during vertical movements, compared to horizontal ones. More specifically, during movement phases when gravity effects can supplement muscle force, muscle activations should be reduced and less positively correlated with movement kinematics. In the present study, we vary movement speed – thus acceleration and deceleration peaks – to further test the *Effort-Optimization Hypothesis* and investigate the relationship between tri-phasic EMG patterns and arm kinematics during vertical arm reaching movements.

## Methods

### Participants

Eleven healthy young volunteers [all males; mean age = 24. 6 ± 0.9 (SD) years; mean height = 178 ± 6 cm; mean weight = 75 ± 8 kg] were included in this study after giving their written informed consent. Participants had normal or corrected-to-normal vision and did not present any neurological or muscular disorders. The study was carried out following legal requirements and international norms (Declaration of Helsinki, 1964) and approved by the French National ethics committee (2019-A01558-49).

### Experimental protocol

Our experimental protocol was similar to those of previous studies investigating gravity-related arm movement control (Gentili et al., 2007; Le Seac’h and McIntyre, 2007; Gaveau et al., 2021). Participants were asked to perform unilateral single-degree-of-freedom vertical and horizontal arm movements (rotation around the shoulder joint) in the sagittal and transversal planes (crossing the right shoulder joint), respectively, at different speeds (i.e. fast, natural and slow). Subjects carried out horizontal movements while being 90° tilted in roll, thereby keeping movement orientation constant between conditions in a body-centered frame of reference (Fig.1A). We investigated single-degree-of-freedom arm movements to isolate and emphasize the mechanical effects of gravity on arm motion. During such movements, only the effect of gravity changes with movement direction. A mechanical system supported the arm during horizontal movements to fully compensating the effects of gravity.

**Fig.1:**
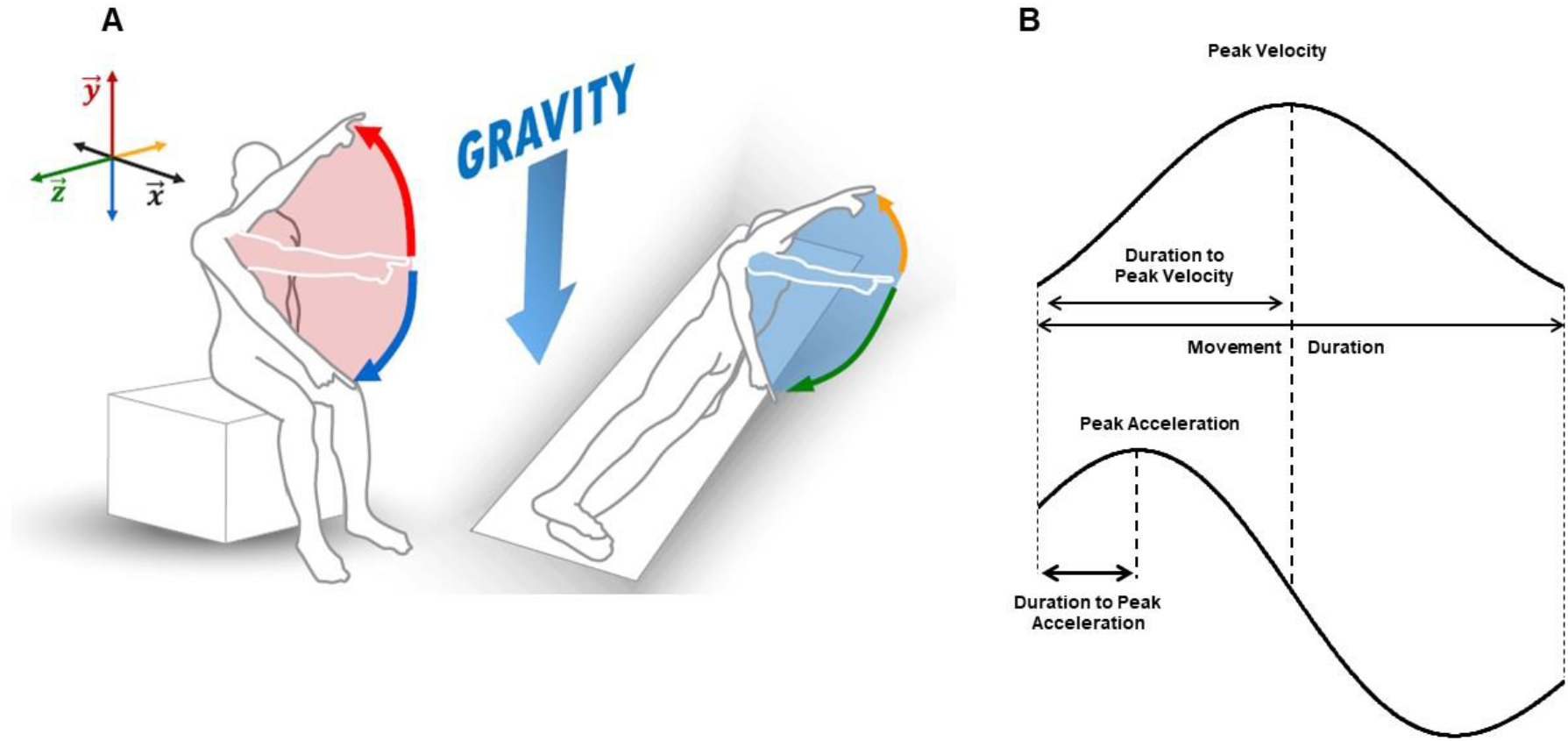
**(A).** Experimental setup. Participants performed arm-pointing movements between two targets, toward the head and feet, in an earth-vertical and an earth-horizontal orientation. Colored arrows depict four different movements directions. **(B).** Illustration of the parameters computed on the velocity and acceleration profiles

The order of orientations was randomized between participants. For each orientation, participants performed movements at three different speeds (fast, natural and slow). Movement speed was also randomized in a block design (three blocks for each orientation). A metronome was used before each block to guide participants. This metronome was set at 150, 100, and 75 bpm for fast, natural and slow blocks respectively. Directions (upward/downward or rightward/leftward) were randomly interleaved within each block. Each participant carried out 360 trials (30 trials x 2 directions x 3 speeds x 2 orientations).

We used a 3D virtual reality environment to set-up the experimental task (HTC Vive; custom environment programed with Unity 3D). Two targets were positioned in a parasagittal plane crossing the participant’s right shoulder joint (parasagittal plane in a body-centered reference frame), at a distance corresponding to the length of their fully extended arm. The targets were symmetrically positioned above and below the antero-posterior horizontal line crossing the shoulder joint. The required angular shoulder rotation between the two targets was 40°, corresponding to a 110° (upward target) and 70° (downward target) shoulder elevation (in a body-centered reference frame).

Each trial was carried out as follows: the experimenter indicated the starting target (red for upward/blue for downward) and the participant positioned his arm fully extended in front of it (initial position). After a brief delay (~2 seconds), the experimenter verbally informed the participant that she/he was free to aim at the other target, as accurately as possible, whenever she/he wanted. Note that reaction time was not emphasized in our experiment. Participants were requested to maintain their final position for a brief period (about 2 seconds) until the experimenter instructed them to relax their arm near the hips. A short rest period (~10 s) separated trials to prevent muscle fatigue. Additionally, a 5mn time interval separated each block. Participants were allowed to perform few practice trials (~5 trials) before each block.

Five reflective markers were placed on the participant’s shoulder (acromion), arm (middle of the humeral bone), elbow (lateral epicondyle), wrist (cubitus styloid process), and finger (nail of the index). We also placed a marker on each target to measure end-point error. Position was recorded with an optoelectronic motion capture system (Vicon system, Oxford, UK; six cameras) at a sampling frequency of 100Hz. The spatial variable error of the system was less than 0.5mm. Additionally, we recorded EMG activity with bipolar surface electrodes (Aurion, Zerowire EMG, sampling frequency: 1000Hz) on the antigravity muscles (anterior deltoid, biceps brachii and the upper head of the pectoralis major) and gravity muscles (posterior deltoid, latissimus dorsi and teres major). The Giganet unit (Vicon, Oxford, UK) recorded kinematic and EMG signals synchronously.

### Data analysis

We processed Kinematic and EMG data using custom programs written in Matlab (Mathworks, Natick, NA). Data processing was similar to previous studies (Poirier et al., 2020; Gaveau et al., 2021).

#### Kinematics analysis

First, we filtered position using a third-order low-pass Butterworth filter (5Hz cut-off, zero-phase distortion, “butter” and “filtfilt” functions). We then computed velocity and acceleration profiles by numerically differentiating (3 points derivative) position signals. Angular joints displacements were computed to ensure that rotations, apart from the shoulder joint, were negligible (<1° for each trial). We defined movement onset and offset as the moments when finger velocity respectively rose above or fell below a threshold corresponding to 10% of peak velocity. Movements presenting multiple local maxima on the velocity profile were automatically rejected from further analysis (< 1% of all trials). The following parameters were then computed from angular profiles of the virtual segment linking finger to shoulder markers (Fig.1B): 1) Movement duration (MD), defined as the duration between movement onset and offset. 2) Peak acceleration (PA), defined as the maximal value of the acceleration profile. 3) Relative duration to peak acceleration (rD-PA), defined as the duration to peak acceleration normalized by total movement duration. 4) Relative duration to peak velocity (rD-PV), defined as the duration to peak velocity normalized by total movement duration.

Additionally, we calculated shoulder inertial torque (SIT) and shoulder gravitational torque (SGT) throughout upward and downward movements using the following equations:

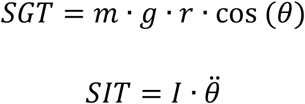

where, *m* is the mass of the arm, *r* is the distance between the center of rotation of the shoulder joint and the center of mass of the whole arm (upper arm + forearm + hand); *g* is the gravitational acceleration (9.81 m·s^-2^); θ represents the angle between the arm and the horizontal axis; *I* is the moment of inertia of the arm around the shoulder joint and 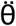 is the angular acceleration of the arm. The values for *m, r* and *I* were calculated for each participant using the anthropometrical data given by Winter (1990).

We obtained a SGT and a SIT profile for each movement. Comparing SIT and SGT, two specific cases could be observed: i) there was a period during acceleration phase where SIT was higher than SGT (Fig. 2A); ii) SIT was systematically smaller than SGT throughout the movement (Fig. 2B). We quantified this phenomenon by subtracting SGT from SIT signal (see Fig.2C and 2D). The resulting profile allowed us to classified trials into two categories: i) SIT > SGT; trials in which the SIT - SGT signal was positive for at least one frame and ii) SIT < SGT; trials in which the SIT - SGT signal was negative during the entire movement. Additionally, for each trial, we computed a parameter quantifying how much SIT exceeded SGT (or conversely). To this aim, we integrated peak torque, defined as the integrated SIT - SGT signal from 50ms before to 50ms after the peak torque (see grey area on Fig.2C and Fig.2D).

**Fig.2:**
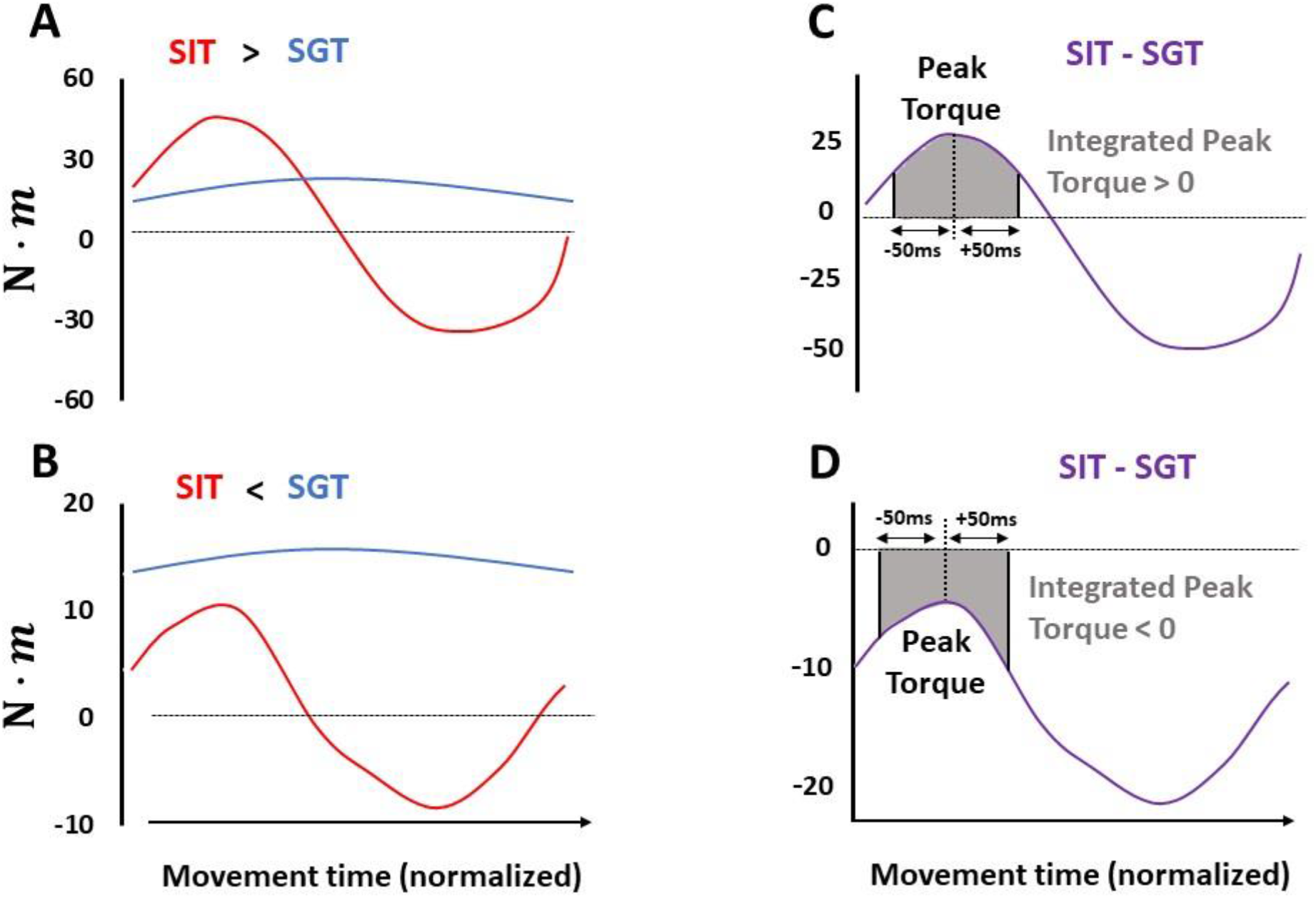
**(A).** Illustration of shoulder inertial torque (SIT) and shoulder gravitational torque (SGT) when SIT > SGT during acceleration phase. **(B).** Illustration of SIT and SGT when SIT < SGT throughout the movement. C, D. Illustration of the SIT-SGT signal (purple line) in each case. Integrated peak Torque is computed as the sum of the signal from 50ms before to 50ms after signal’s maxima.

#### EMG analysis

EMG signals were first filtered using a bandpass third-order Butterworth filter (bandpass 20-300Hz, zero-phase distortion, “butter” and “filtfilt” functions) then rectified. Signals were integrated using a 50ms sliding window and cut off from 250ms before movement onset to 250ms after movement offset. Then, signals were filtered using a low-pass third-order Butterworth filter (low-pass frequency: 20Hz) and normalized by the maximal EMG value recorded during maximal isometric voluntary contractions (MIVC). At the beginning of the experiment, participants performed three flexions and three extensions MIVC at a shoulder angle of 90°. Lastly, EMG traces were classified according to movement mean velocity and averaged across three trials (from the three slowest to the three fastest movements), resulting in 10 EMG traces to be analyzed for each block. Each set of three traces was normalized in duration (corresponding to the mean duration of the three traces) before averaging.

We computed the phasic component of each EMG signal using a well-known subtraction procedure (Flanders and Herrmann, 1992; Buneo et al., 1994; Flanders et al., 1994; D’Avella et al., 2006, 2008; Gaveau et al., 2021). We averaged the values of the integrated EMG signals from 500ms to 250ms before movement onset and from 250ms to 500ms after movement offset (Fig.3A). The tonic component was calculated as a linear interpolation between these two averaged values (Fig.3B). Finally, we computed the phasic component by subtracting the tonic component from the full EMG signal (Fig.3C).

**Fig.3:**
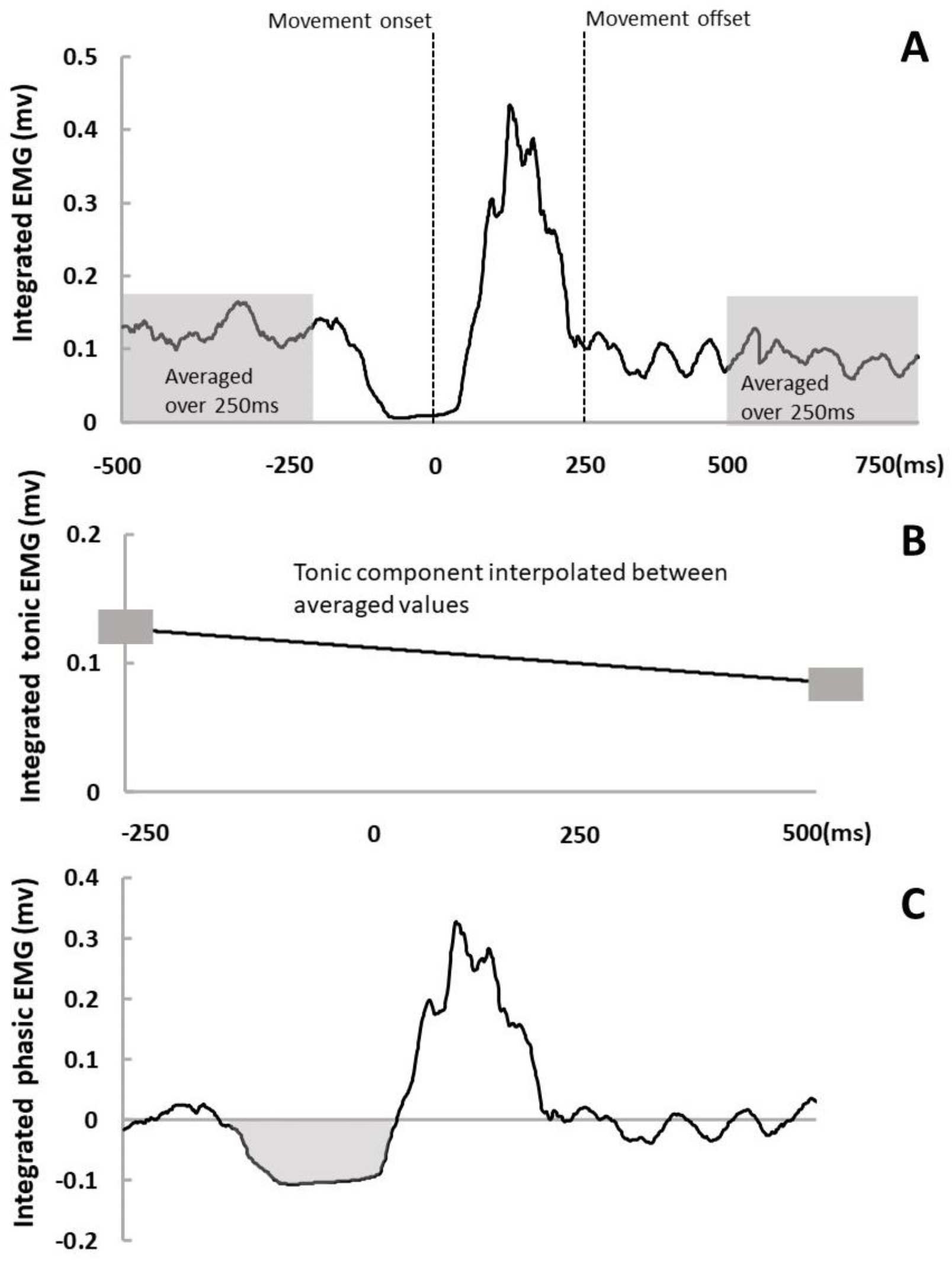
Analysis of the tonic and phasic components of the EMG. **(A).** Illustration of the integrated full EMG signal. Movement onset and offset were detected on the velocity profile. We averaged the signal from 500 to 250ms before movement onset and from 250 to500ms after movement offset (grey windows). **(B).** Tonic EMG component was obtained by linear interpolation between the two previously averaged values (small grey rectangles). **(C).** Phasic EMG component was obtained by subtracting the tonic component (in panel B) from the full EMG (in panel A).

Following the report of negative epochs in phasic EMG signal of antigravity muscles during vertical movements (Gaveau et al., 2021; Poirier et al., 2022), we also quantified the negativity of phasic signals in antigravity muscles for each condition. We computed integrated negative values over acceleration and deceleration phases, and normalized it by the respective phase duration to account for differences in movement duration.

Additionally, for each trial, we computed the following parameters. 1) Integrated agonist signal, defined as the integrated phasic signal over acceleration phase normalized by acceleration duration. 2) Integrated antagonist signal, defined as the integrated phasic signal over deceleration phase normalized by deceleration duration. Antigravity muscles were defined as agonist muscles for upward and leftwards (= towards head) movements and as antagonist muscles for downwards and rightwards (= towards feet) movements. Gravity muscles were defined as agonist muscles for downwards and rightwards (= towards feet) movements and as antagonist muscles for upward and leftwards (= towards head) movements.

#### Statistics

Data were analyzed using repeated measures variance analysis (three-way ANOVA) with 3 factors: *speed* (fast vs natural vs slow), *body orientation* (upright vs tilted), and *direction* (head vs feet). *HSD-Tukey tests* were used for post-hoc comparisons. We analyzed correlations between variables with repeated measures correlations (rmcorr package in R, Bakdash and Marusich, 2017). The level of significance was set at p<0.05 for all analyses.

## Results

In this experiment, we asked eleven subjects to perform one degree of freedom horizontal and vertical movements around the shoulder joint at different speeds (i.e. fast, natural and slow). Subjects carried out horizontal movements while being 90° tilted in roll, thereby keeping movement orientation constant between conditions in the body-centered frame of reference (see Fig. 1A). Thus, the same muscles were involved in producing vertical and horizontal movements. We used a mechanical system to support horizontal movements and fully compensate gravity effects. We recorded arm kinematics and EMG activity of antigravity (anterior deltoid, pectoralis major upper head, and biceps brachii) and gravity muscles (Posterior Deltoid, Latissimus Dorsi and Teres major). In this section, we only present statistical results that relate to our main hypothesis. To provide a more exhaustive statistical description of the results, values for all parameters and statistical effects are presented in Table.1 & 2.

**Table.1:** Kinematic parameters. Mean (±SD) movement duration (MD), angular amplitude (Amp), peak acceleration (PA), relative duration to peak acceleration (rD-PA) and relative duration to peak velocity (rD-PV) in each body orientation and direction (Upright: upward and downward. Tilted: leftward (toward head) and rightward (toward feet) for each speed (Fast, Natural and Slow).

|  | Fast |  |  |  | Natural |  |  |  | Slow |  |  |  |
| --- | --- | --- | --- | --- | --- | --- | --- | --- | --- | --- | --- | --- |
|  | Upright |  | Tilted |  | Upright |  | Tilted |  | Upright |  | Tilted |  |
|  | Up | Down | Left (Head) | Right (Feet) | Up | Down | Left (Head) | Right (Feet) | Up | Down | Left (Head) | Right (Feet) |
| **MD (ms)** | 409 ± 77 | 412 ± 81 | 498 ± 98 | 484 ± 91 | 604 ± 133 | 623 ± 125 | 691 ± 198 | 705 ± 198 | 901 ± 266 | 937 ± 248 | 1002 ± 356 | 1007 ± 335 |
| **Amp (°)** | 41.4 ± 2.8 | 40.2 ± 2.9 | 41.1 ± 3.6 | 42.1 ± 3.7 | 41.0 ± 2.6 | 40.3 ± 2.4 | 41.1 ± 3.2 | 41.5 ± 2.9 | 40.4 ± 2.4 | 39.9 ± 2.4 | 40.6 ± 3.1 | 40.6 ± 3.1 |
| **PA (m/s<sup>2</sup>)** | 19.4 ± 5.4 | 18.5 ± 6.7 | 12.8 ± 4.9 | 12.9 ± 4.9 | 8.5 ± 2.7 | 6.7 ± 2.0 | 6.4 ± 2.2 | 5.9 ± 2.2 | 5.0 ± 2.6 | 3.9 ± 1.4 | 3.6 ± 1.5 | 3.2 ± 1.4 |
| **rD-PA** | 0.20 ± 0.02 | 0.25 ± 0.04 | 0.20 ± 0.03 | 0.22 ± 0.03 | 0.18 ± 0.03 | 0.22 ± 0.05 | 0.18 ± 0.03 | 0.17 ± 0.03 | 0.14 ± 0.04 | 0.17 ± 0.04 | 0.17 ± 0.04 | 0.15 ± 0.04 |
| **rD-PV** | 0.45 ± 0.02 | 0.48 ± 0.04 | 0.45 ± 0.04 | 0.48 ± 0.03 | 0.44 ± 0.04 | 0.50 ± 0.04 | 0.47 ± 0.03 | 0.48 ± 0.03 | 0.42 ± 0.04 | 0.46 ± 0.04 | 0.44 ± 0.06 | 0.47 ± 0.03 |

**Table.2:** F and P values of all main ANOVA effects are presented for all parameters. Red fonts highlight significant main effects. Those statistics are presented for descriptive purposes only. MD: Movement Duration; rD-PA: Relative Duration to Peak Acceleration; rD-PV: Relative Duration to Peak Velocity; PA: Peak Acceleration; Neg Acc: Integrated Negativity on Acceleration phase; Neg Dec: Integrated Negativity on Deceleration phase; Int Ag Acc: Integrated Agonist Activity on Acceleration phase; Int Ant Dec: Integrated Antagonist Activity on Deceleration phase.

| Effect | Speed |  | Direction |  | Orientation |  | Speed x Orientation |  | Speed x Direction |  | Orientation x Direction |  | Speed x Orientation x Direction |  |
| --- | --- | --- | --- | --- | --- | --- | --- | --- | --- | --- | --- | --- | --- | --- |
|  | F | P | F | P | F | P | F | P | F | P | F | P | F | P |
| MD | 45.69 | 1<sup>-6</sup> | 2.34 | 0.16 | 8.37 | 0.02 | 0.01 | 0.99 | 2.45 | 0.11 | 4.47 | 0.06 | 1.14 | 0.34 |
| rD-PA | 37.79 | 1<sup>-6</sup> | 14.93 | 3.1<sup>-3</sup> | 5.20 | 4.5<sup>-2</sup> | 3.27 | 0.06 | 16.10 | 6.8<sup>-5</sup> | 47.47 | 4.2<sup>-5</sup> | 0.93 | 0.41 |
| rD-PV | 3.88 | 0.04 | 23.89 | 6.4<sup>-4</sup> | 0.83 | 0.38 | 2.55 | 0.10 | 0.07 | 0.93 | 7.34 | 0.02 | 3.33 | 0.06 |
| PA | 59.17 | 4.0<sup>-9</sup> | 13.81 | 4.0<sup>-3</sup> | 36.31 | 1.3<sup>-4</sup> | 25.50 | 1<sup>-6</sup> | 0.44 | 0.65 | 6.37 | 0.03 | 0.34 | 0.10 |
| Neg Acc | 16.87 | 5.1<sup>-5</sup> | 58.98 | 1.5<sup>-5</sup> | 50.03 | 3.4<sup>-5</sup> | 13.91 | 1.6<sup>-4</sup> | 16.44 | 1.0<sup>-5</sup> | 57.32 | 1.9<sup>-5</sup> | 14.73 | 1.2<sup>-4</sup> |
| Neg Dec | 7.11 | 4.6<sup>-3</sup> | 12.37 | 5.6<sup>-3</sup> | 24.92 | 5.4<sup>-4</sup> | 5.33 | 0.01 | 12.22 | 3.4<sup>-4</sup> | 8.21 | 0.02 | 8.99 | 1.6<sup>-3</sup> |
| Int Ag Acc | 27.40 | 2.0<sup>-6</sup> | 60.65 | 1.5<sup>-5</sup> | 0.07 | 0.79 | 6.3<sup>-4</sup> | 0.99 | 28.14 | 2.0<sup>-6</sup> | 55.72 | 2.1<sup>-5</sup> | 6.30 | 7.6<sup>-3</sup> |
| Int Ant Dec | 15.17 | 9.8<sup>-5</sup> | 17.37 | 1.9<sup>-3</sup> | 0.14 | 0.72 | 3.02 | 0.07 | 22.82 | 7.0<sup>-6</sup> | 1.70 | 0.22 | 4.39 | 0.03 |

### Kinematics

First, we briefly present kinematic results to test whether they comply with the *Effort-Optimization Hypothesis*. Previous work has revealed directional asymmetries in the vertical plane such that relative durations to peak acceleration (rD-PA) and velocity (rD-PV) are shorter and peak acceleration (PA) are larger for upward than for downward movements (Gentili et al., 2007; Le Seac’h and McIntyre, 2007; Gaveau et al., 2011; 2014; 2016; 2021; Yamomoto et al., 2014; 2016; 2019; Rousseau et al., 2016; Poirier et al., 2020; Hondzinski et al., 2016; Bringoux et al., 2012). These specific asymmetries are thought to represent the kinematic signature of an *Effort-Optimization* process that optimizes gravity effects to minimize muscle effort (Berret et al., 2008; Crevecoeur et al., 2009; Gaveau et al., 2014, 2016, 2021). The present results well align with previous observations. All three parameters revealed a significant interaction effect between movement direction (head or feet) and body orientation (vertical or horizontal): rD-PA, F = 47.47, p = 4.2e-5, ⊓_p_^2^ = 0.83; rD-PV, F = 7.34, p = 0.02, ⊓_p_^2^ = 0.42; PA, F = 6.37, p = 0.03, ⊓_p_^2^ = 0.39. As previously observed, rD-PA and rD-PV were shorter and PA were larger for upward than for downward movements (see Fig.4 and Table.1) No such effect or a significantly reduced one existed in the horizontal plane.

**Fig.4:**
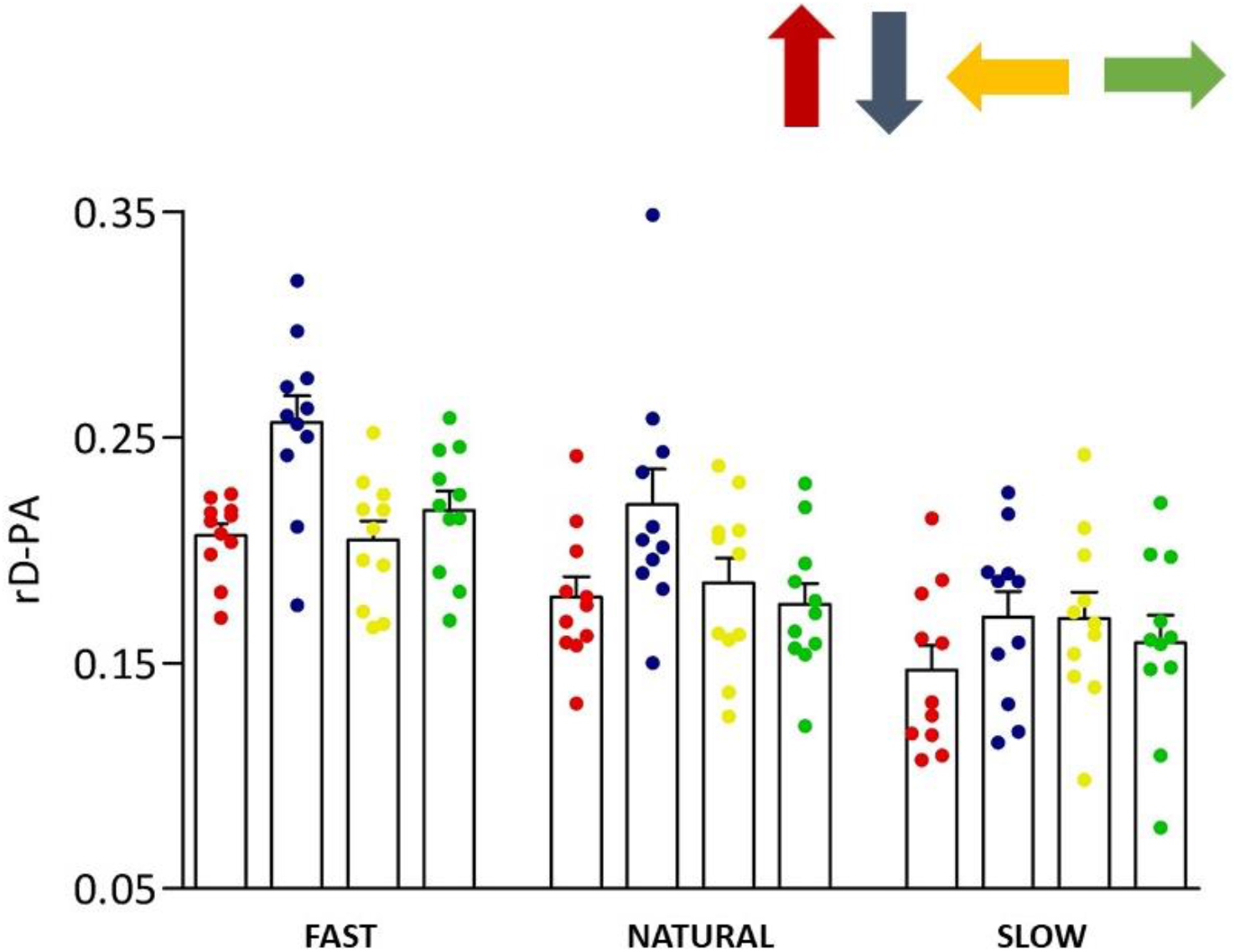
Mean (±SE) relative duration to peak acceleration (rD-PA) in each direction for fast, natural, and slow movements. Each dot represents a participant.

### EMG

We computed phasic EMG signal following a well-known method (see Methods, Fig.3 and Flanders and Herrman, 1992; D’Avella et al., 2008; Gaveau et al. 2021; Poirier et al. 2022). According to the compensation hypothesis, EMG signals can be decomposed into two components; i) the tonic corresponding to the muscle force that compensates gravity effects throughout the motion; ii) the phasic component, corresponding to the muscle force that changes velocity. If muscle force indeed compensates for gravity effects, throughout the motion, the phasic component should remain superior or equal to zero. Contrariwise, the *Effort-Optimisation Hypothesis* proposes that the CNS uses gravity effects to discount muscle effort. In other words, gravity should assist in accelerating downward and decelerating upward movements. Consequently, the amount of muscular activation during these two phases should decrease under the theoretical level needed to compensate for gravity torque (i.e., below zero in a phasic signal), highlighting the assisting role of gravity for muscular effort minimization. Supporting the *Effort-Optimization Hypothesis*, Gaveau et al. (2021) recently reported consistent negative epochs in arm flexor muscles during vertical movements. Furthermore, those negative epochs were specific to movement’s direction: they occurred during the acceleration phase of downward movements and during the deceleration phase of upward movements. For each movement speed and direction, Fig.5 presents the mean phasic EMG profiles recorded in the present study. The present results qualitatively reproduce those of Gaveau et al. (2021) and extend them to new movement speeds (natural and slow).

**Fig.5:**
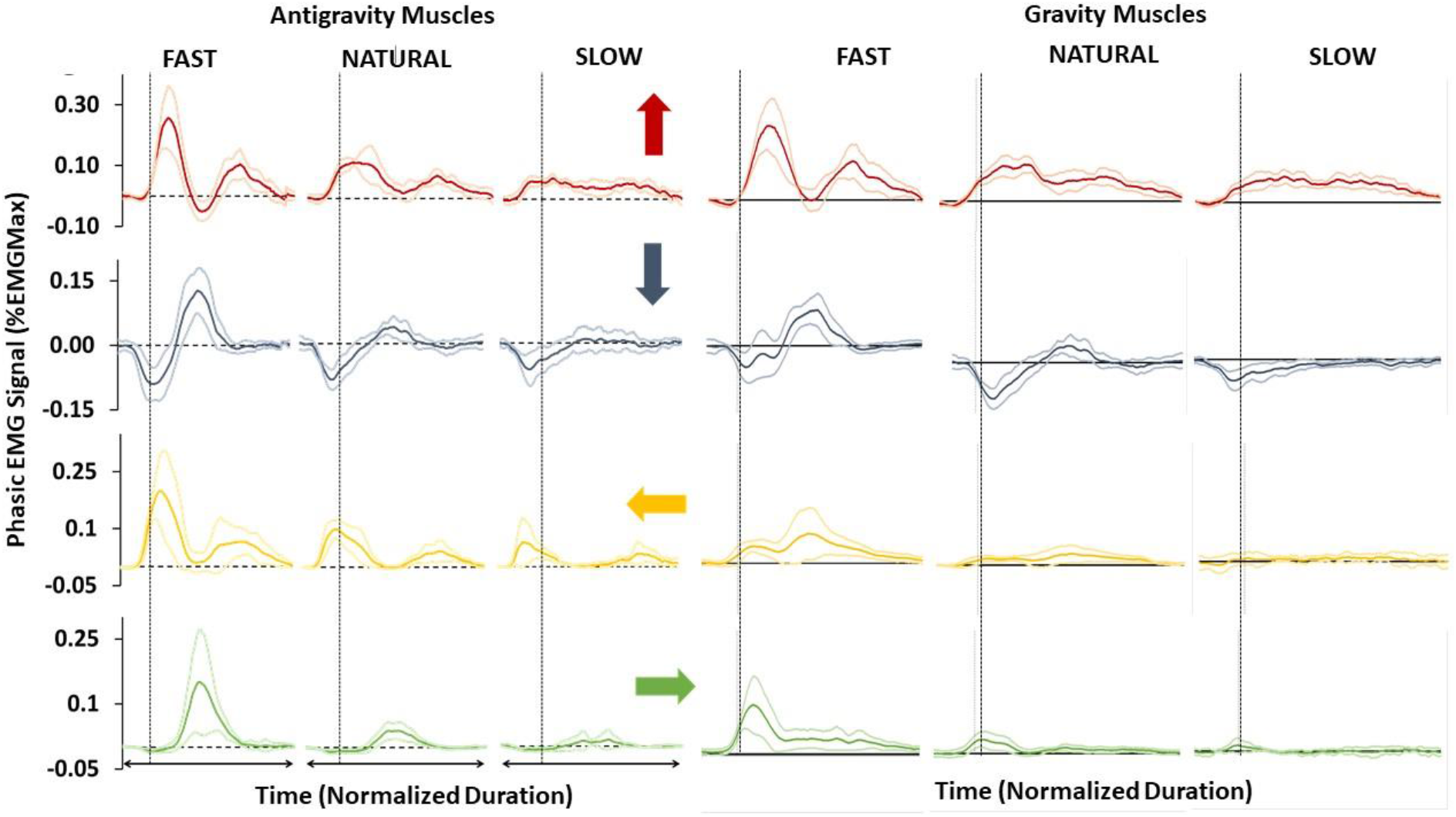
Average phasic signal (mean±SD) for arm flexor (muscles pulling toward head, left panel) and extensor (muscles pulling toward feet, right panel) in each direction for each speed. The dashed line represents the kinematic onset of the movement.

### Antigravity muscles Negativity

We first investigated whether movement speed influences gravity effects optimization by quantifying antigravity muscles deactivation below the theoretical tonic level. We integrated the amount of negativity for all directions and speeds, during the acceleration (Fig.6A) and the deceleration phases (Fig.6B). For each trial, we normalized the integrated negativity by phase duration. During the acceleration phase (Fig.6A), we observed a significant interaction effect between speed x orientation x direction (F = 14.73; p = 1.2e-4; ⊓_p_^2^ = 0.60). For all three speeds, negativity was significantly higher for downward movement compared to all other directions (p < 1.6-e-4; Cohen’s d > 1.36 for all comparisons), supporting the specific exploitation of gravity effects for the acceleration of downward movements. Negativity in downward movements also progressively decreased with movement speed (p < 0.01; Cohen’s d > 0.99). There were no significant differences between other directions (p > 0.27; Cohen’s d < 1.12 for all comparisons). During the deceleration phase (Fig.6B), the interaction effect between speed x orientation x direction also reached significance (F = 8.99; p = 1.6e-3; ⊓_p_^2^ = 0.47). HSD-Tukey test revealed that negativity was higher for upward movements, compared to other movement directions, at the fast speed only (p < 1.6e-4; Cohen’s d > 1.03 for each comparison; See Fig.6B). This supports the specific exploitation of gravity effects for the deceleration of upward movements. Similarly to downward direction during the acceleration phase, the amount of negativity during deceleration of upwards movements qualitatively decreased with movement speed (see Fig.6B; post-hoc comparisons for the upward condition: fast vs natural: p = 6.3e-4; Cohen’s d = 0.80; fast vs slow: p = 1.6e-4; Cohen’s d = 0.94; natural vs slow: p = 0.86; Cohen’s d =0.96). There was no significant difference between other directions at natural and slow speeds (p > 0.11; Cohen’s d < 1.30 in each case).

**Fig.6:**
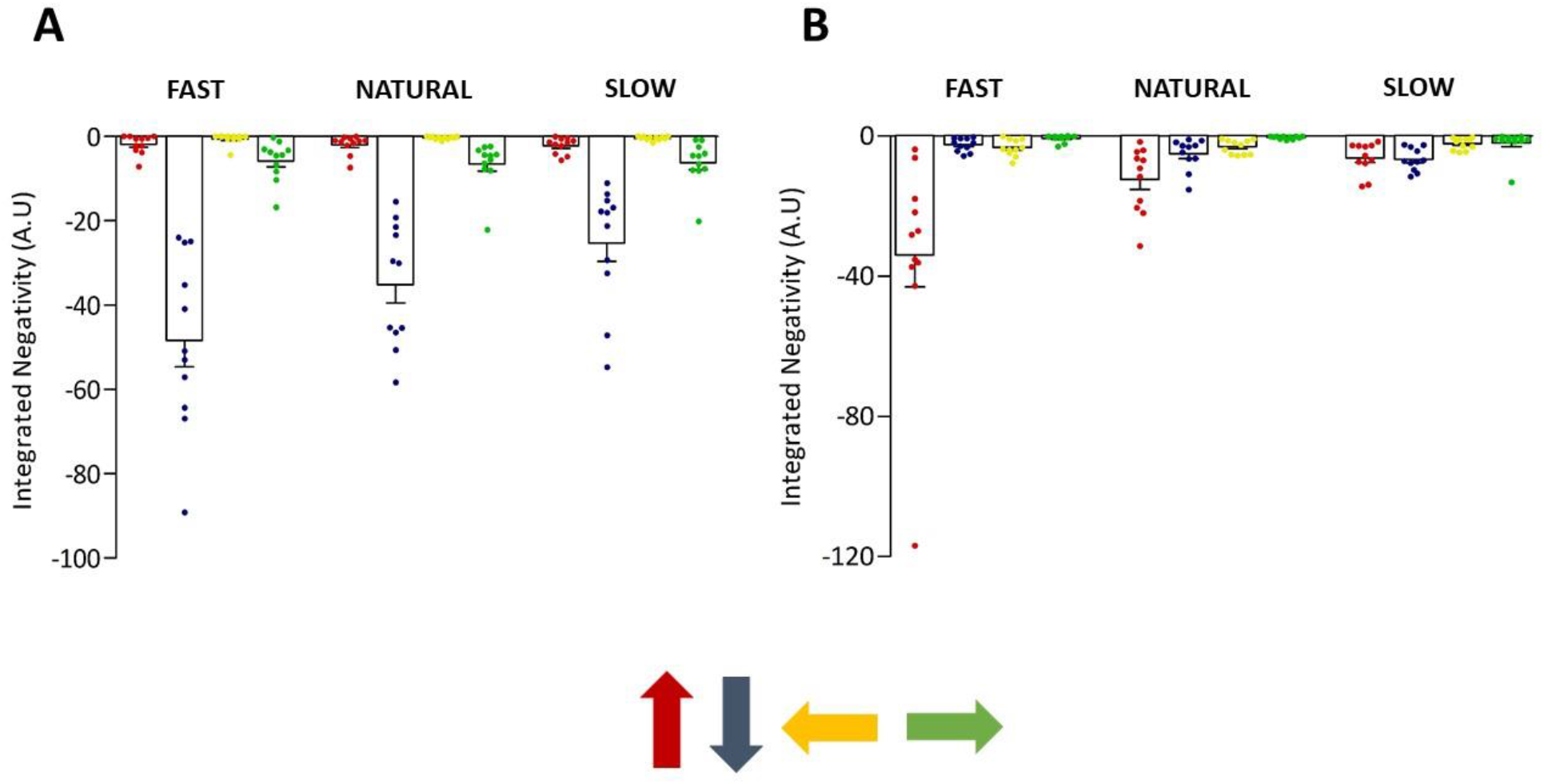
**(A).** Mean (±SE) integrated negativity during acceleration phase in each direction for fast, natural, and slow movements. Each dot represents a participant. **(B).** Mean (±SE) integrated negativity during deceleration phase in each direction for fast, natural, and slow movements. Each dot represents a participant.

These results add to recent ones supporting the *Effort-Optimization Hypothesis* (Gaveau et al., 2021; Poirier et al., 2022). They reveal that movement speed influences how much the CNS harvests gravity effects to move the arm. More they suggest that this speed effect differs between upward and downward movements.

### Gravity muscles activation

Next, we investigate whether antigravity muscle deactivations allow discounting the effort spend by gravity muscles; i.e. muscles working in the same sense as gravity. Due to the full body tilt we operated between vertical and horizontal movements, the same muscles acted towards the head or feet in the two planes (sagittal and transverse). This allows us to perform within-muscle comparisons between vertical and horizontal movements to compare gravity muscles effort between the two planes. We integrated phasic signals of gravity muscles during the acceleration phase of movements directed towards the feet (Fig.7A) and during the deceleration phase of movements directed towards the head (Fig.7B); i.e. when negativity of antigravity muscles was observed. For each trial, integrated signals were normalized by the length of the corresponding movement phase duration (in ms). According to the *Effort-Optimization Hypothesis*, we expected activations to be smaller for downward compared to rightward (toward feet) movements, during the acceleration phase, and for upward compared to leftward (toward head) movements, during the deceleration phase.

**Fig.7:**
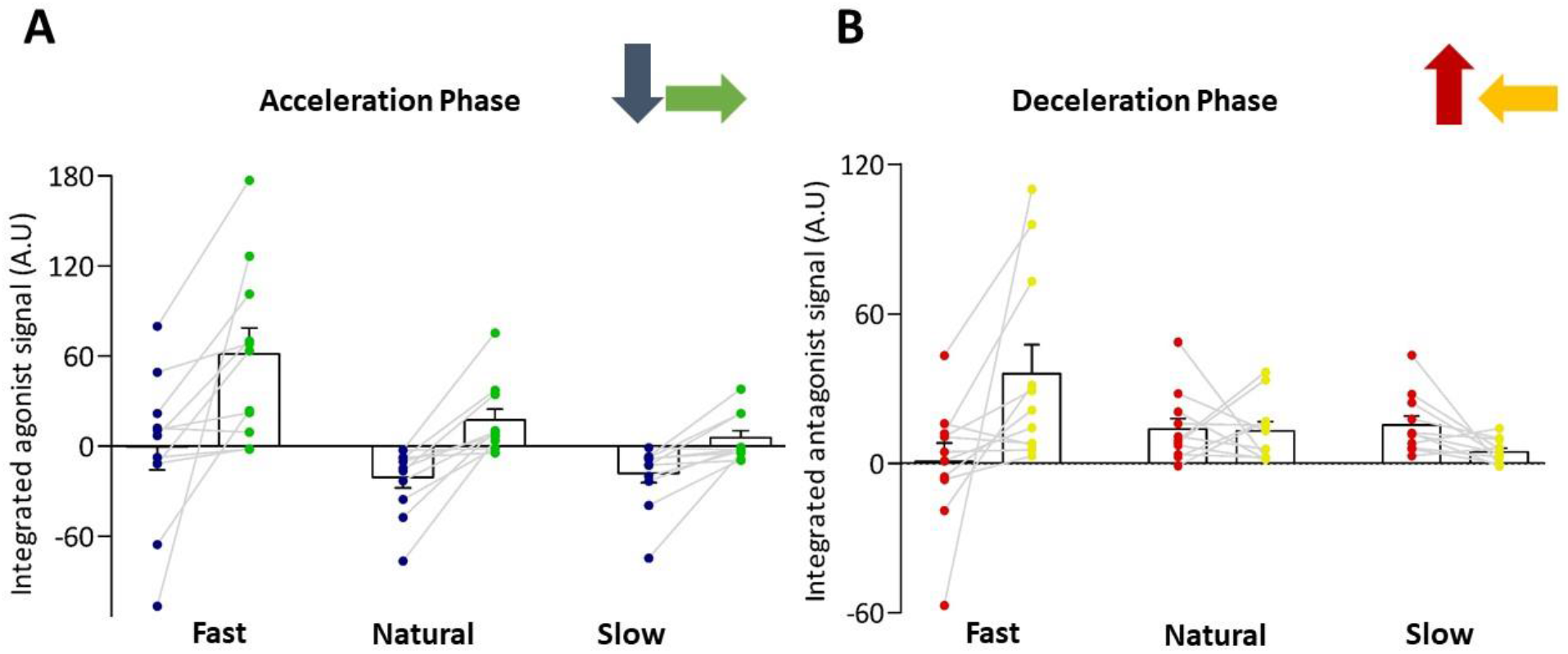
**(A).** Mean (±SE) gravity muscles integrated phasic signal during acceleration phase in downward (blue dots) and leftward (green dots) movements for each speed. **(B).** Mean (±SE) gravity muscles integrated phasic signal during deceleration phase in upward (red dots) and rightward (yellow dots) movements for each speed. Each dot represents a participant.

During the acceleration phase, the interaction effect between speed x orientation x direction significantly influenced muscle activations (F = 6.30; p = 7.6e-3; ⊓_p_^2^ = 0.39; see Fig.7A). Gravity muscles were significantly less activated during downward than during rightward movements at fast speed (p = 8e-4; Cohen’s d = 0.80). Although this effect did not reach significance at natural and slow speeds, comparisons revealed large to very large effect sizes (natural, p = 0.07, Cohen’s d = 1.45; slow, p = 0.59, Cohen’s d = 1.17). During the deceleration phase, the interaction effect between speed x orientation x direction also significantly influenced muscle activations (F = 4.36; p = 0.03 ⊓_p_^2^ = 0.30; see Fig.7B). Gravity muscles were significantly less activated for upward than for leftward movements at fast speed only (p = 0.04; Cohen’s d = 0.85). At natural and slow speeds, comparisons led to high p values and small effect sizes (Natural, p = 1.00, Cohen’s d = 0.05; Slow, p = 0.99, Cohen’s d = 0.91).

Gravity muscles activation confirm the beneficial effect of antigravity muscles deactivation to discount the effort spent by gravity muscles. Similarly to results on antigravity muscles negativity, gravity muscles activations reveal an effect of movement speed on this effort-discounting process. Again, the results suggest that this speed effect differs between upward and downward movements.

### Dynamic analysis

To try and explain the speed effects observed on the effort-discounting process, we performed a dynamic analysis. Our rationale was that the speed effect may be explained by the balance between inertial and gravitational torques. For a fast downward movement, inhibiting antigravity muscles during the acceleration phase may not be sufficient to produce the net force that will move the arm at the required speed. However, for a slow downward movement, gravity effects alone may be sufficient to accelerate the arm. Conversely, for an upward movement, gravity effects may help decelerating the arm if the instant velocity is high enough (if the movement is fast). If the instant velocity is low, deactivating antigravity muscles below the tonic level – i.e. the level required to compensate gravity torque – is not possible as it would quickly lead to a reversal of the arm movement sense. This simple rationale suggests that the balance between inertial and gravity torques may produce a direction-dependent effect on the effort discounting process. In other words, this process would be favored by movement slowness during downward movements whilst it would be favored by movement rapidness during upward movements. We hypothesized that, during the deceleration phase of upward movements, gravity muscles would be less activated when the Shoulder Inertial Torque (SIT) exceeds the Shoulder Gravity Torque (SGT) than when SIT < SGT. Inversely, during the acceleration phase, we hypothesized that gravity muscles would be less activated when SIT < SGT than when SIT > SGT.

During the acceleration phase, SIT peaked above SGT in 178 out of 330 upward movements and in 144 out of 330 downward movements (see Methods and Fig.2). To finely test the dependence of gravity muscles activity on the balance between SIT and SGT, we computed a parameter quantifying how much SIT peaked above or below SGT during the acceleration of each movement. We called it *Integrated Peak Torque*. The higher SIT compared to SGT, the higher the value of Integrated Peak Torque. Using repeated measure correlations, we found that Integrated Peak Torque (computed during the acceleration phase) negatively correlated with the deceleration-related gravity muscles’ activity of upward movements (r_rm(317)_ = −0.25; IC95% = [−0.352; −0.146]; p < 4.95e-6, see Fig.8A). Thus, faster movements took greater advantage of gravity effects when moving upwards. During downward movements, repeated measure correlations revealed that Integrated Peak Torque positively correlated with the acceleration-related gravity muscles’ activity (r_rm(317)_ = 0.50; IC95% = [0.410; 0.577]; p < 1e-6; see Fig.8B). Thus, slower movements took greater advantage of gravity effects when moving downwards.

**Fig.8:**
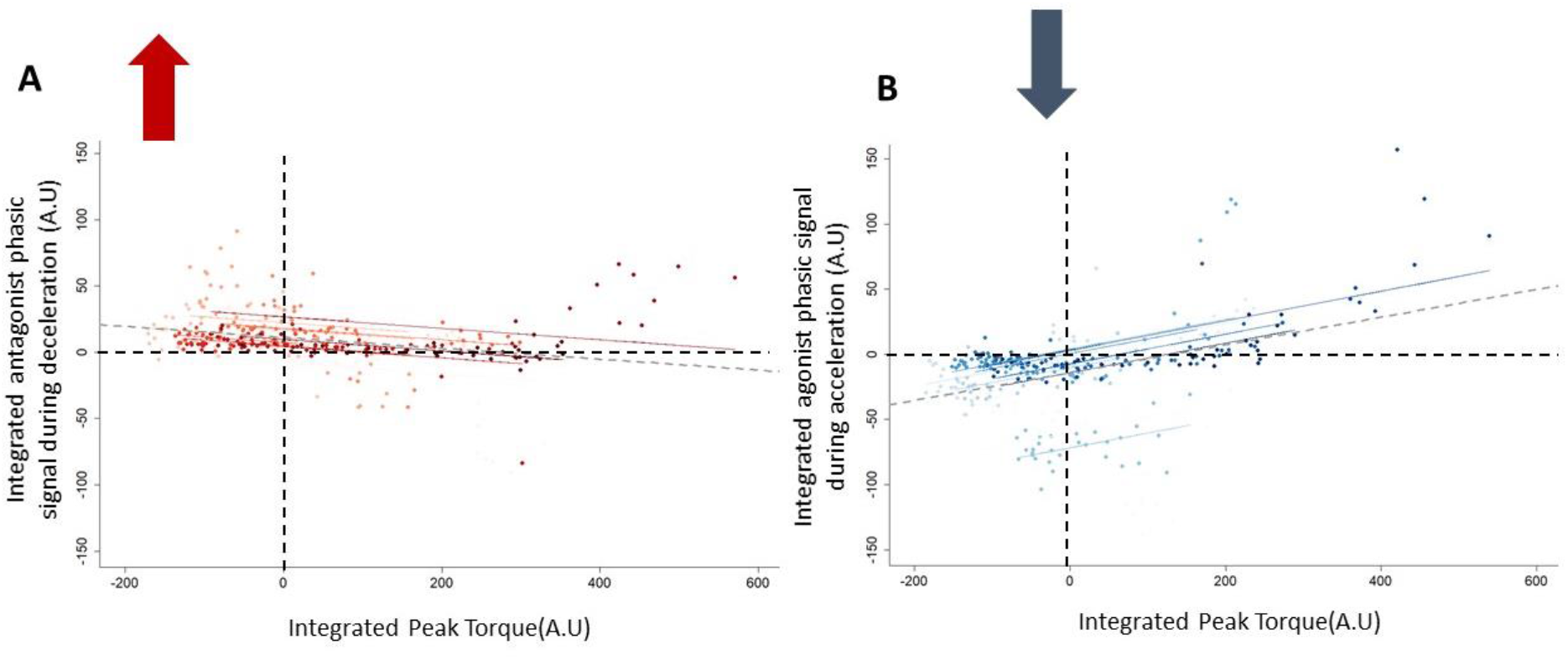
**(A).** Repeated measures correlation between Integrated Peak Torque and integrated antagonist phasic signal during deceleration phase. **(B).** Repeated measures correlation between Integrated Peak Torque and integrated agonist phasic signal during acceleration phase. Each dot represents a trial and each color represents a participant.

This simple analysis of movement dynamics and EMG patterns supports the hypothesis that, at least in part, pure environmental physics explains the direction-dependent effect of speed on the effort-discounting process. This process is favored by movement slowness during downward movements and by movement rapidness during upward movements.

### EMG to kinematics relationship

Last, we investigate how the effort discounting process affects the well-known relationship between muscle activation and movement kinematics. The literature on the tri-phasic patterns (for a review see Berardelli et al., 1996) has consistently shown that, during the acceleration and deceleration phases of horizontal movements, the integrated signal of agonist (muscles pulling toward the target) and antagonist muscles (those pulling away from the target) positively correlates with peak acceleration and deceleration respectively (Corcos et al., 1989; Gottlieb et al., 1989a; Cooke and Brown, 1994). Here we used repeated measure correlations to probe the EMG to kinematics relationship across vertical and horizontal movements.

Fig.9 presents the relationships between peak acceleration and agonist muscle activation during the acceleration phase in all four directions. We obtained significant correlations for upward (r_rm(317)_ = 0.93, IC95% = [0.914, 0.944], p < 1e-6; see Fig.9A), downwards (r_rm(317)_ = 0.50, IC95% = [0.416, 0.582], p < 1e-6; See Fig. 9B), leftward (r_rm(317)_ = 0.85, IC95% = [0.811, 0.875], p < 1e-6; see Fig.9C) and rightward movements (r_rm(317)_ = 0.81, IC95% = [0.773, 0.848], p < 1e-6; see Fig.9D). Considering 95% confidence intervals on rrm - values, it can be observed that 95%CI overlapped between rightwards and leftwards, but that those of upwards and downwards stood alone. The 95%CI of upward and downward movements were respectively superior and inferior to those of horizontal movements.

**Fig.9.**
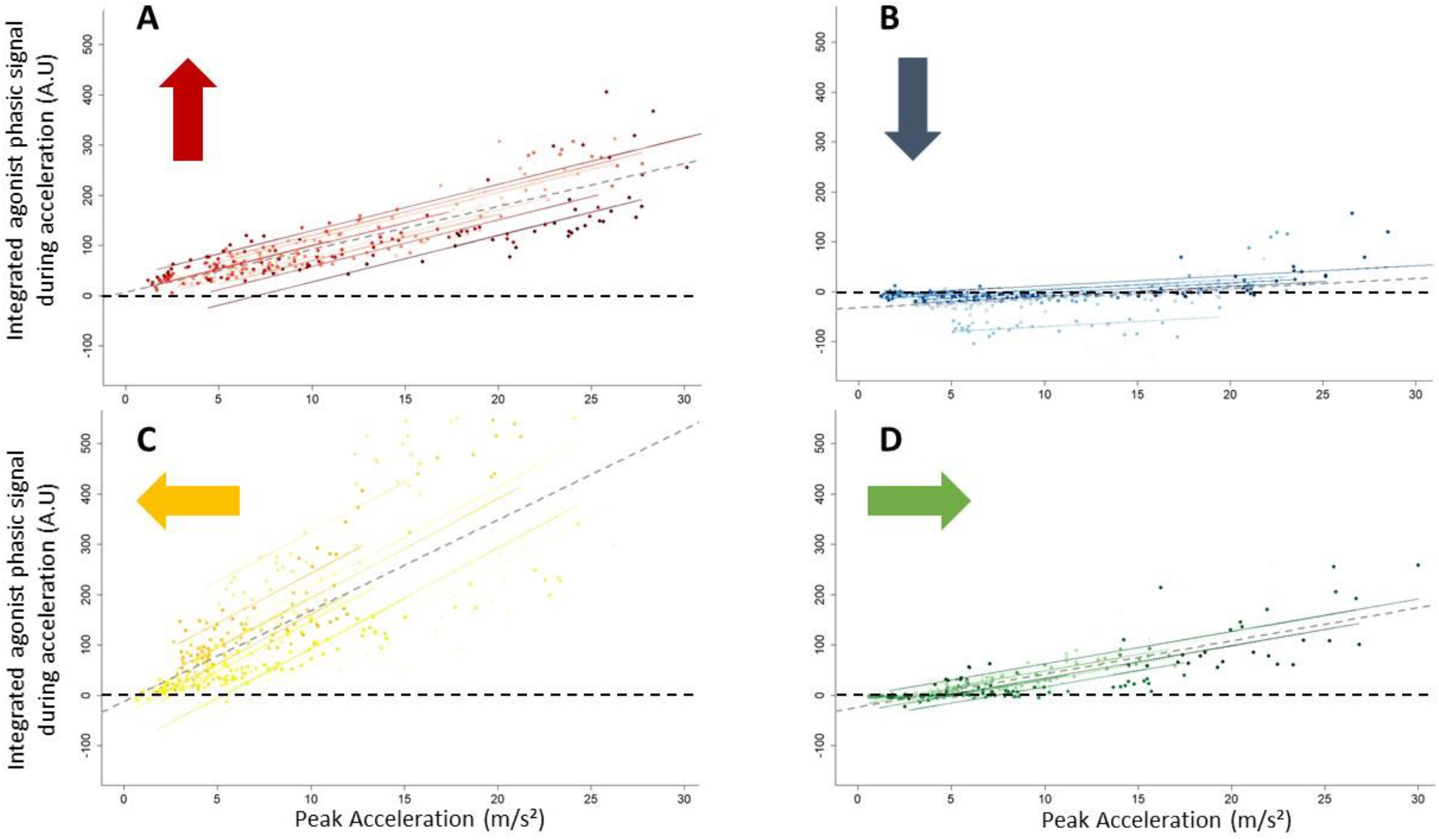
Repeated measures correlation between peak acceleration and integrated agonist phasic signal during acceleration phase for upward **(A)**, downward **(B)**, leftward **(C)**, and rightward **(D)** movements. Each dot represents a trial and each color represents a participant. The axis scale is kept constant between panels to ease comparisons between directions. The grey dashed line represents the mean regression for all participants.

Fig.10 presents the relationships between peak deceleration and antagonist muscle activation during the deceleration phase in all four directions. We observed significant positive correlations for downward (r_rm(317)_ = 0.89, IC95% = [0.862, 0.909], p < 1e-6; see Fig.10B), leftward (r_rm(317)_ = 0.73, IC95% = [0.676, 0.779], p < 1e-6; see Fig.10C), and rightward movements (r_rm(317)_ = 0.67, IC95% = [0.605, 0.727], p < 1e-6; see Fig.10D). For upward movements, however, we found that integrated antagonist activation negatively correlated with peak deceleration (r_rm(317)_ = −0.29; IC95% = [−0.386; −0.184]; p < 1e-6; see Fig.10A). Again, although the 95%CI of rightward and leftward movements overlapped, those of upward and downward movements stood alone.

**Fig.10:**
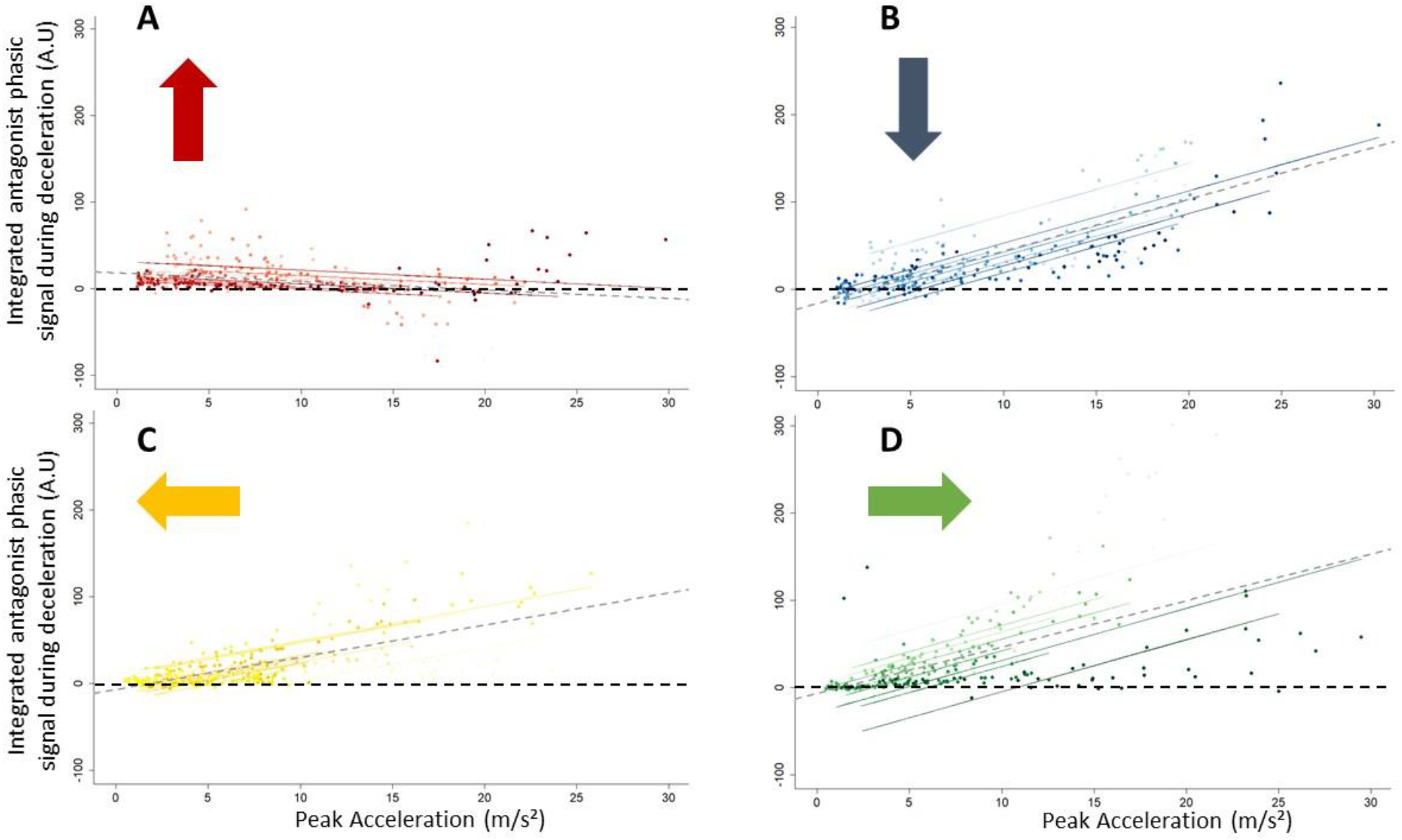
Repeated measures correlation between peak deceleration and integrated antagonist phasic signal during deceleration phase for upward **(A)**, downward **(B)**, leftward **(C)**, and rightward **(D)** movements. Each dot represents a trial and each color represents a participant. The axis scale is kept constant between panels to ease comparisons between directions. The grey dashed line represents the mean regression for all participants.

Overall, these correlational analyses reveal that the EMG to kinematics relationship differ between vertical and horizontal movements. They even more strongly differ between upward and downward movements. During phases when gravity assists muscle force, EMG to kinematics relationships are particularly altered compared to horizontal movements. These alterations further support the rationale according to which the effort-discounting process is favored by movement slowness when mowing downwards and by movement rapidness when moving upward.

## Discussion

In this study we recorded arm kinematics and electromyographic activity of antigravity and gravity muscles during fast, natural and slow arm reaching movements performed in a vertical and a horizontal orientation. We quantified phasic negative activity of antigravity muscles for all directions and found that this negativity was more important during the acceleration phase of downward movements and during the deceleration phase of upward movements, resulting in diminished phasic activity compared to horizontal movements. Furthermore, we found direction-specific effects of movement speed on phasic EMG activity of gravity muscles. This resulted in altered EMG to kinematics relationships in vertical movements compared to horizontal ones. Those results support the *Effort-minimization hypothesis* and confirm that the negativity of phasic EMG is an important aspect of the motor command. More, the present results reveal that the CNS finely tunes this feature across a range of movement speeds and directions.

Consistently with previous results (Gentili et al., 2007; Le Seac’h and McIntyre, 2007; Gaveau et al., 2011; 2014; 2016; 2021; Yamomoto et al., 2014; 2016; 2019; Rousseau et al., 2016; Poirier et al., 2020; Hondzinski et al., 2016; Bringoux et al., 2012), we found direction-dependent kinematics during vertical movements. Specifically, rD-PA, rD-PV were found to be shorter for upward than for downward movements. Additionally, peak acceleration was higher for upward than for downward movements. Those directional asymmetries were absent (or significantly reduced) in horizontal movements, as testified by the *orientation x direction* effect observed for all these parameters. These results further support that direction-dependent kinematics are the hallmark of the gravity-related effort optimization strategy.

As previously observed (Gaveau et al., 2021; Poirier et al. 2022), we also found substantial negativity in phasic activity of the antigravity muscles for vertical movements only. This negativity appeared specifically in the acceleration phase of downward movements and during the deceleration phase of upward movements. Negativity in the phasic muscle activity implies that this activity is inferior to the theoretical activity necessary for compensate for gravity torque. Thus, this negativity observed in specific phases of vertical movements (i.e when gravity is coherent with the arm’s acceleration) is thought to be the neural signature of a motor optimization process that exploits gravity force to discount muscle force (Gaveau et al., 2021). This is corroborated by the reduced phasic muscular activity of gravity muscles (the muscles pulling in the same sense as gravity) in vertical compared with horizontal movements (see Fig.7B and 7C).

The novelty of our work concerns the quantification of the effects of movement speed on the effort discounting process. In the acceleration phase, whatever the movement speed, the negativity of phasic antigravity muscles EMGs was significantly higher in downward compared to other movements directions. However, during the deceleration phase, the negativity of phasic antigravity muscles EMGs was more important in the upward direction, at the fast speed only. We hypothesized that the effort discounting process depended on the simple balance between inertial and gravity torques. Specifically, we thought that the CNS could not exploit gravity effects during an upward movement when the inertial torque was inferior to the gravitational torque. In this case, deactivating antigravity muscles beyond the level necessary to compensate gravity torque would compromise the integrity of the movement (gravity would pull the arm down and the movement would reverse direction). Thus, upward movements must be performed at a certain minimum speed for the CNS to exploit gravity effects in replacing the force that would otherwise be produced by gravity muscles. During a downward movement, conversely, the CNS can exploit gravity effects to produce movement acceleration at slow speeds. When the speed requirement increases, the CNS must recruit gravity muscle to add the supplementary force that is required to accelerate the arm. Overall, this rationale postulates that gravity-related effort-optimization would be favored by movement rapidness when moving upwards and by movement slowness when moving downwards. This hypothesis was confirmed by our dynamical analysis. We computed a parameter quantifying whether the shoulder inertial torque (SIT) peaked above or below the shoulder gravitational torque (SGT). We found that the integrated activity of antigravity muscles negatively and positively correlated with this parameter during upward and downward movements respectively.

From experiments investigating the control of horizontal movements, it was previously thought that the muscle activity responsible for arm acceleration/deceleration scaled according to peak acceleration/deceleration (Corcos et al., 1989; Gottlieb et al., 1989; Cooke and Brown 1994). Here, the integrated antagonist phasic activity during the deceleration phase was negatively correlated with peak deceleration for upward movements, whereas this correlation was positive for downward and horizontal movements (Fig.10). The integrated agonist phasic activity during the acceleration phase was positively correlated with peak acceleration for downward movements, but less so than in other movement directions (Fig.9).

These results indicate that, because of the role of gravity in the production of arm motion, the scaling of muscle activity according to inertial forces is not generalizable to vertical arm movements. Furthermore, this suggests that fast movements could be more energetically-optimal than slow movements during upward movements. This is consistent with the kinematic result showing that directional asymmetries are more important for fast movements (Gaveau and Papaxanthis, 2011).

Collectively, our results extend previous findings supporting the *Effort-optimization hypothesis*. We demonstrated that the phasic muscular activity during vertical movements is subtly tuned to movement speed and direction. Investigating this gravity-related tuning may represent a valuable tool to probe motor control in various condition.

